# Primary cilia formation does not rely on WNT/β-catenin signaling

**DOI:** 10.1101/2020.10.30.361642

**Authors:** Ondrej Bernatik, Petra Paclikova, Anna Kotrbova, Vitezslav Bryja, Lukas Cajanek

**Affiliations:** Masaryk University, Faculty of Medicine, Dept. of Histology and Embryology, Kamenice 5, Brno 62500, Czech Republic; Masaryk University, Faculty of Science, Dept. of Experimental Biology, Section of Animal Physiology and Immunology, Kamenice 5, Brno 62500, Czech Republic

## Abstract

Primary cilia act as crucial regulators of embryo development and tissue homeostasis. They are instrumental for modulation of several signaling pathways, including Hedgehog, WNT, and TGF-β. However gaps exist in our understanding of how cilia formation and function is regulated.

Recent work has implicated WNT/β-catenin signaling pathway in the regulation of ciliogenesis, yet the results are conflicting. One model suggests that WNT/β-catenin signaling negatively regulates cilia formation, possibly via effects on cell cycle. In contrast second model proposes a positive role of WNT/β-catenin signaling on cilia formation, mediated by the re-arrangement of centriolar satellites in response to phosphorylation of the key component of WNT/β-catenin pathway, β-catenin.

To clarify these discrepancies, we investigated possible regulation of primary cilia by the WNT/β-catenin pathway in cell lines (RPE-1, NIH3T3, HEK293) commonly used to study ciliogenesis. We used WNT3a to activate or LGK974 to block the pathway, and examined initiation of ciliogenesis, cilium length, and percentage of ciliated cells. We show that the treatment by WNT3a has no- or lesser inhibitory effect on cilia formation. Importantly, the inhibition of secretion of endogenous WNT ligands using LGK974 blocks WNT signaling but does not affect ciliogenesis. Finally, using knock-out cells for key WNT pathway components, namely DVL1/2/3, LRP5/6 or AXIN1/2 we show that neither activation nor deactivation of the WNT/β-catenin pathway affects the process of ciliogenesis.

These results suggest that WNT/β-catenin-mediated signaling is not generally required for efficient cilia formation. In fact, activation of the WNT/β-catenin pathway in some systems seems to moderately suppress ciliogenesis.

## Introduction

Primary cilia are tubulin-based rod-shaped organelles on the surface of most mammalian cells. They play a fundamental role in embryo development and tissue homeostasis. Importantly, defects in primary cilia structure and function lead to variety of developmental disorders collectively called ciliopathies (Mitchison and Valente 2017; Reiter and Leroux 2017; Hildebrandt, Benzing, and Katsanis 2011). Moreover, primary cilia defects have been related to cancer (Wong et al. 2009; Han et al. 2009; Jenks et al. 2018).

Cilium formation is organized by the mother centriole (MC)-derived basal body, the older centriole of the pair that makes up the centrosome. While centrosome is best known as microtubule organizing center coordinating mitosis, primary cilium formation is tightly connected with G1/G0 phase (Mirvis, Stearns, and Nelson 2018; Ford et al. 2018). The growth of primary cilium itself is preceded by the accumulation of vesicles at MC distal appendages (Sorokin 1962; Westlake et al. 2011; Schmidt et al. 2012; Lu et al. 2015; Wu, Chen, and Tang 2018) and by the removal of CEP97/CP110 capping complex specifically from MC distal end (Spektor et al. 2007). Major role in the cilia initiation is linked to the Tau tubulin kinase 2 (TTBK2) activity (Goetz, Liem, and Anderson 2012). Once recruited to MC by distal appendage protein CEP164 (Čajánek and Nigg, 2014; Oda *et al.,* 2014), TTBK2 seems to control both the process of vesicle docking and the CP110/CEP97 removal (Goetz, Liem, and Anderson 2012; Lo et al. 2019). In turn, this allows the extension of tubulin-based axoneme sheathed by ciliary membrane from MC-derived basal body. The formed cilium is physically separated from the rest of a cell by ciliary transition zone, a selective barrier ensuring only specific proteins to enter the cilium (Gonçalves and Pelletier 2017; Nachury 2018; Garcia-Gonzalo and Reiter 2017). Such compartmentation and hence specific protein composition of primary cilium is the basis for its instrumental role in the Hedgehog signaling pathway in vertebrates (Bangs and Anderson 2017; Nachury and Mick 2019). In addition, several links between primary cilia and other signaling pathways such as WNT or TGF-β have recently emerged (Anvarian et al. 2019).

WNT signaling pathways are developmentally important signaling routes regulating cell differentiation, migration and proliferation and their activity controls shaping of the embryo (Nusse and Clevers 2017). WNT signaling pathways can be distinguished based on whether they use β-catenin as an effector protein. The pathway relying on stabilization of β-catenin is termed the WNT/β-catenin pathway and regulates stemness, cell differentiation and proliferation, while the β-catenin-independent or noncanonical WNT pathways regulate cytoskeleton, cell polarity and cell movements (Steinhart and Angers 2018; Humphries and Mlodzik 2018). These two branches of WNT pathways are activated by a distinct set of extracellularly secreted WNT ligand proteins (Angers and Moon 2009). WNTs are posttranslationally palmitoylated by O-Acyl-transferase Porcupine, and only after the lipid modification are the WNT proteins fully active (Zhai, Chaturvedi, and Cumberledge 2004; Willert et al. 2003). Following their secretion, WNTs bind to seven-pass transmembrane receptors from Frizzled family that form heterodimeric complexes with various coreceptors. WNT/β-catenin pathway uses LRP5/6 coreceptors (Tamai et al. 2000; Wehrli et al. 2000; Pinson et al. 2000). Signal received by the receptor-coreceptor pair on the cell membrane is then relayed to Dishevelled (DVL) proteins that are used both by the noncanonical and the WNT/β-catenin pathways (Sokol 1996; Wallingford et al. 2000). β-catenin destruction complex, composed of proteins Adenomatous polyposis coli (APC), AXIN and two kinases; GSK3-β and CK1-α, is then inactivated by DVL sequestration of AXIN proteins (Tamai et al. 2004). Then β-catenin phosphorylation by GSK3-β and CK1-α on its N-terminal degron is terminated and the nonphosphorylated Active β-catenin (ABC) accumulates, translocates to the nucleus where it binds transcription factors of TCF-LEF family to trigger transcription of target genes (Molenaar et al. 1996; Behrens et al. 1996). Not surprisingly, many developmental disorders and cancers are directly caused by WNT pathways deregulation (Zhan, Rindtorff, and Boutros 2017; Humphries and Mlodzik 2018).

Whilst the connections between primary cilia and hedgehog signaling are well documented (Huangfu et al. 2003; Corbit et al. 2005; Rohatgi, Milenkovic, and Scott 2007), the relationship between cilia and WNT signaling is rather controversial. The exception here seems to be the WNT/PCP pathway (one of the noncanonical WNT pathways (Butler and Wallingford 2017)), which was described to affect cilia formation and functions via effects on cytoskeleton and basal body positioning (Wallingford and Mitchell 2011; May-Simera and Kelley 2012; Carvajal-Gonzalez, Mulero-Navarro, and Mlodzik 2016; Bryja, Červenka, and Čajánek 2017). As for the WNT/β-catenin pathway, there are reports showing that primary cilia loss or disruption leads to upregulation of the pathway activity (Corbit et al. 2008; Lancaster, Schroth, and Gleeson 2011; Zingg et al. 2018; B. Liu et al. 2014; McDermott et al. 2010; Wiens et al. 2010) but also studies that deny any involvement of primary cilia in WNT/β-catenin signaling (Ocbina, Tuson, and Anderson 2009; Huang and Schier 2009). Some of these discrepancies can perhaps be explained by the effects of ciliary components directly on WNT/β-catenin pathway, independently of their role in cilia formation, as has been recently showed for several regulators of ciliary transport (M. Kim et al. 2016; Balmer et al. 2015).

To make the matters even more puzzling, two opposing models have recently emerged regarding possible function of WNT/β-catenin pathway in cilia formation. Activation of the WNT/β-catenin pathway in neural progenitors of the developing cerebral cortex was reported to hamper cilia formation in mice (Nakagawa et al. 2017), arguing for a negative role of the excesive WNT/β-catenin signaling in ciliogenesis. In contrast, a recent report described a direct involvement of WNT/β-catenin signaling pathway in promotion of primary cilia formation through β-catenin driven stabilization of centriolar satellites in RPE-1 cell line (Kyun et al. 2020). We approached this conundrum using cell lines that commonly serve as ciliogenesis model systems (RPE-1, NIH3T3, HEK293). Using either pharmacological or genetic means to manipulate the WNT/β-catenin pathway, we found no evidence of facilitated ciliogenesis in response to the activation of WNT/β-catenin signaling.

## Material and Methods

### Cell culture

RPE-1 cells were grown in DMEM/F12 (ThermoFisher Scientific, 11320033) supplemented by 10% FBS (Biosera, cat. No. FB-1101/500), 1% Penicillin/Streptomycin (Biosera, cat. No. XC-A4122/100) and 1% L-glutamine (Biosera, cat. No. XC-T1715/100), HEK293 T-Rex (referred to as HEK293, cat.no. R71007, Invitrogen) and NIH3T3 cells were grown in DMEM Glutamax^®^ (Thermo Fisher Scientific, 10569069) supplemented by 10% FBS and 1% Penicillin/Streptomycin. RPE-1 cells were starved by serum free medium, NIH3T3 cells were starved by 0.1% FBS containing medium. Cells were seeded at 50000/well (RPE-1 and NIH3T3) or 120000/well (HEK293) of 24 well plate. Treatments by small molecules were done for indicated times: LGK974 (0.4 μM) (Sellcheck, cat. No. S7143) for 72h (LGK974 was re-added to the starvation medium as indicated in Fig. 2A), Cytochalasin D (500nM) (Merck Cat. No. C8273) for 16h. WNT3a (90 ng/ml) (R&D systems, Cat.no. 5036-WN) for 2 or 24h.

### Western Blot, quantification

WB was performed as previously described (Bernatik et al. 2020). Antibodies used: LRP6 (Cell signaling, Cat.no. #2560), Phospho-LRP5/6 (Ser1493/Ser1490; Cell signaling, Cat.no. #2568), AXIN1 (Cell signaling, Cat.no. #3323) DVL2 (Cell signaling, Cat.no. #3216), Active-β-catenin (Merck, Cat. no. 05-665-25UG), α-tubulin (Proteintech, Cat.no. 66031-1-Ig). Quantifications were performed using Fiji distribution of ImageJ. Intensity of ABC band was measured and normalized to mean value from all conditions of given experiment. Intensity of DVL2 was calculated as the ratio of the DVL2 upper to lower band intensity (the bands are indicated by arrows in the corresponding Figures) and normalized to mean value from all conditions of given experiment. Quantification was performed on n=3. Statistical analyses by students t-test or one-way ANOVA were performed using Graphpad Prism, P<0.05 (*), P<0.01 (**), P<0.001 (***).

### Immunocytochemistry

RPE-1 cells were seeded on glass coverslips, treated as indicated, washed by PBS and fixed for 10min in −20°C methanol, washed 3x by PBS, blocked (2% BSA in PBS with 0.01% NaN_3_), 3x washed by PBS, incubated with primary antibodies for 1h, 3x washed by PBS, incubated with secondary antibodies (Goat anti-Rabbit IgG Alexa Fluor 488 Secondary Antibody, Cat.no. A11008; Goat anti-Mouse IgG Alexa Fluor 568 Secondary Antibody, Cat.no. A11031, all from Thermo Fisher Scientific) for 2h in dark, washed 3x by PBS, incubated 5 min with DAPI, 2x washed by PBS and mounted to glycergel (DAKO #C0563). Microscopy analysis was done using Zeiss AxioImager.Z2 with Hamamatsu ORCA Flash 4.0 camera, 63x Apo oil immersion objective, and ZEN Blue 2.6 acquisition SW (Zeiss). Image stacks acquired using Zeiss AxioImager.Z2 were projected as maximal intensity images by using ImageJ distribution FIJI (Schindelin et al. 2012). Where appropriate, contrast and/or brightness of images were adjusted by using Photoshop CS5 (Adobe) or FIJI. To assess effects on ciliogenesis or cilia length, at least 4-5 fields of vision (approximately 200-400 cells per experiment) were analyzed per experimental condition, on at least n=3. Cilia present on HEK293 cells were counted manually. Cilia present on RPE-1 or NIH3T3 were counted in ACDC software semiautomatic mode, all cilia present were verified and adjusted manually as recommended (Lauring et al. 2019). For the experiments in Suppl. Fig. 1A, 1B, 1C, and 1D (analysis of CP110 and TTBK2 presence on the MC), 3-4 fields of vision (200-400 cells) were analyzed per experimental run, n=3. Statistical analyses by one-way ANOVA were performed using Graphpad Prism, P<0.05 (*), P<0.01 (**), P<0.001 (***) and P<0.0001 (****). Results are presented as mean plus SEM. Primary antibodies used: Arl13b (Proteintech, Cat.no. 17711-1-AP), γ-tubulin (Merck, T6557), CP110 (Proteintech, 12780-1-AP), TTBK2 (Merck, Cat.no. HPA018113).

**Figure 1:**
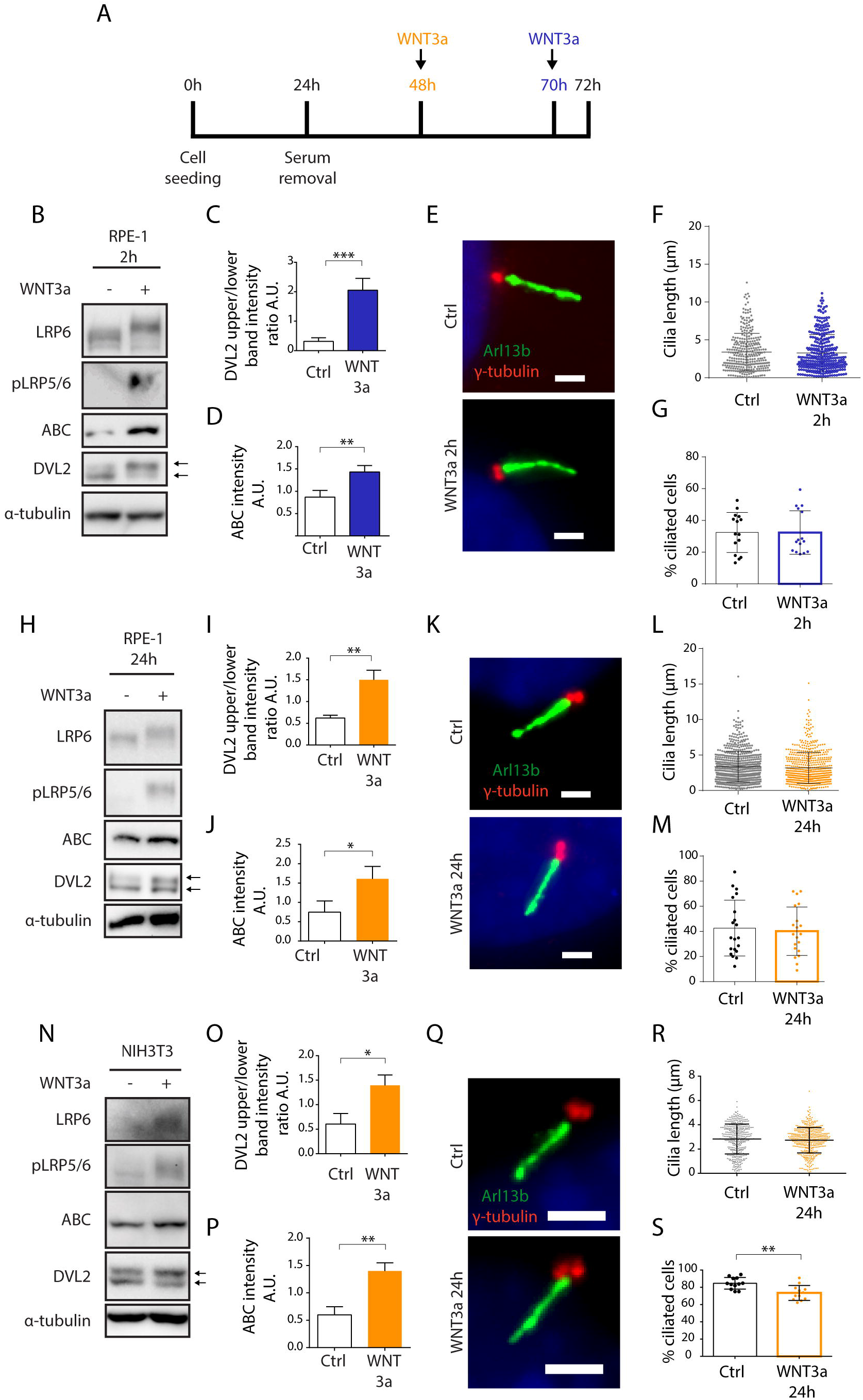
WNT3a does not promote ciliogenesis or cilia length. **(A)** Experimental scheme of WNT3a treatment experiment. Cells were seeded and grown for 24h, then starved for additional 48h. 2h treatment (RPE-1) by WNT3a is indicated in blue, 24h treatment is indicated in orange (RPE-1 and NIH3T3). **(B)** Western blot analysis of 2h WNT3a treatment of RPE-1. The treatment leads to LRP6 shift and increased LRP5/6 phosphorylation, DVL2 phosphorylation and upshift, and accumulation of ABC. The quantitation of DVL2 band intensities (upper to lower band intensity ratio, the bands are indicated by arrows) is shown in **(C)** n=3, the quantification of relative ABC levels is presented in **(D)** n=3. **(E)** Representative images of RPE-1 cells treated by WNT3a or vehicle (control) for 2h and stained for Arl13b (green) and γ-tubulin (red). Scale bar 2μm. DAPI (blue) was used to counter stain nuclei. The corresponding quantification of the cilia length **(F)** and the percentage of cells with Arl13+ cilium **(G).** Each dot indicates either length of a single primary cilium (F) or percentage of ciliated cells in a single image (G). **(H)** Western blot analysis of 24h WNT3a treatment of RPE-1. The treatment leads to LRP6 shift, increased LRP5/6 phosphorylation, DVL2 phosphorylation and upshift, and accumulation of ABC. Quantification of DVL2 bands (indicated by arrows) intensity ratio is shown in **(I)** n=3, quantification of relative ABC levels is presented in **(J)** n=3**. (K)** Representative images of RPE-1 cells treated by WNT3a or vehicle (control) for 24h and stained for Arl13b (green) and γ-tubulin (red). Scale bar 2μm. DAPI (blue) was used to counter stain nuclei. The corresponding quantification of the cilia length and the percentage of cells with Arl13+ cilium is shown in **(L)** and **(M)**, respectively. Each dot indicates either length of a single primary cilium (L) or percentage of ciliated cells in a single image (M) n=4. **(N)** Western blot analysis of NIH3T3 cells treated by WNT3a for 24h shows LRP6 shift and LRP5/6 phosphorylation, DVL2 phosphorylation and upshift, and accumulation of ABC. Quantification of DVL2 band intensities (upper to lower band intensity ratio, the bands are indicated by arrows) is shown in **(O)** n=3, quantification of relative ABC intensity **(P)** n=3. **(Q)** Representative images of NIH3T3 cells treated by WNT3a for 24h, stained for Arl13b (green), and γ-tubulin (red). Scale bar 2μm. DAPI (blue) was used to counter stain nuclei. The corresponding quantification of the cilia length **(R)** and the percentage of cells with Arl13+ cilium **(S).** Each dot indicates either length of a single primary cilium (R) or percentage of ciliated cells in one image frame (S) n=3.

### Dual luciferase (TopFLASH) assay, transfection of HEK293

Transfection and dual luciferase assay of HEK293 WT and KO cells was carried out as previously described (Paclíková et al. 2017). In brief, in 0.1 μg of the pRLtkLuc plasmid and 0.1 μg of the Super8X TopFlash plasmid per well of 24 well plate were cotransfected, on the next day cells were treated by 90ng/ml WNT3a and signal was measured after 24h treatment.

### CRISPR/Cas9 generation of LRP5/6 double knock-out and AXIN1/2 double knock-out HEK293 cells

Used guide RNAs were following: LRP5 gRNA gagcgggccgacaagactag, LRP6 gRNA ttgccttagatccttcaagt, AXIN1 gRNA cgaacttctgaggctccacg, and AXIN2 gRNA tccttattgggcgatcaaga. gRNAs were cloned into pSpCas9(BB)-2A-GFP (PX458) (Addgene plasmid, 41815) or pU6-(BbsI)_CBh-Cas9-T2A-mCherry (Addgene plasmid, 64324) plasmids. Following transfection by Lipofectamine^®^ 2000 (Thermo Fisher Scientific) the transfected cells were FACS sorted (FACSAria Fusion (BD Biosciences)) and clonally expanded. Genotyping of LRP5 KO and AXIN2 KO mutants was done following genomic DNA isolation (DirectPCR Lysis Reagent; 301-C, Viagen Biotech) by PCR using DreamTaq DNA Polymerase (Thermo Fisher Scientific). Used primers: LRP5 forward: gttcggtctgacgcagtaca, LRP5 reversed: aggatggcctcaatgactgt, AXIN2 forward: cagtgccaggggaagaag, AXIN2 reversed: gtcttggtggcaggcttc. PCR products were cut by BfaI (R0568S, NEB) in case of LRP5 KO and Hpy188III (R0622S, NEB) for AXIN2 KO screening, respectively. Successful disruption of individual ORFs was confirmed by sequencing, Suppl. Fig. 2A and 2D.

## Results

### Treatment by recombinant WNT3a induces WNT/β-catenin pathway activation but not ciliogenesis

First, we tested if primary ciliogenesis can be modulated by activation of WNT/β-catenin pathway in RPE-1 by recombinant WNT3a. Experiment outline is schematized (Fig. 1A). We initially treated the cells for 2 hours. While we observed the expected accumulation of active β-catenin (ABC), phosphorylation of LRP5/6 coreceptors (pLRP5/6, S1490/S1493), and phosphorylation and upshift of DVL2 (Fig. 1B, 1C, 1D), the WNT3a did not alter length or number of Arl13b positive cilia (Fig. 1E, 1F, 1G). Next, we examined effects of prolonged treatment of RPE-1 cells by WNT3a. Importantly, we were able to detect that WNT/β-catenin pathway is still active after 24h, as visible from the elevated levels of ABC, pLRP5/6 or DVL2 phosphorylation (Fig. 1H, 1I, 1J), but the treatment did not show any notable effects on cilia length or numbers (Fig. 1K, 1L, 1M). In agreement with these data, WNT3a treatment failed to alter either TTBK2 recruitment to MC (Suppl. Fig. 1A, 1B) or MC-specific loss of CP110, (Suppl. Fig. 1C, 1D). To corroborate these findings, we also tested the influence of WNT3a in NIH3T3 cell line. Similarly to RPE-1, WNT3a treatment for 24h was able to activate the WNT/β-catenin pathway in NIH3T3 cells (Fig. 1N, 1O, 1P), but the length of cilia was not affected, (Fig. 1Q, 1R). Intriguingly, we detected a decrease in the percentage of ciliated cells following the WNT3a treatment (Fig. 1S).

### Inhibition of WNT secretion halts WNT signaling but not ciliogenesis

Having found WNT3a-activated WNT/β-catenin signaling is not sufficient to promote cilia formation, we tested a possibility that steady state WNT signaling is required for effective ciliogenesis. WNT/β-catenin pathway is intensively studied as a driver of oncogenic growth, thus there are currently available various small molecules that inhibit WNT ligand secretion. To this end, we used a Porcupine inhibitor LGK974 to block the secretion of endogenous WNT ligands and in turn block the steady state WNT signaling (Jiang et al. 2013). As a positive control in these experiments we used cytochalasin D (CytoD), an actin polymerization inhibitor known to facilitate ciliogenesis and promote cilia elongation (J. Kim et al. 2015). Experiment outline is schematized (Fig. 2A). As expected, we observed that LGK974 treatment led to downshift of DVL2 (Fig, 2B, 2C) confirming the endogenous WNT signaling was successfully ablated. Importantly however, the LGK974 treatment did not alter primary ciliogenesis, in contrast to CytoD that facilitated CP110 removal from MC (Suppl. Fig 1C, 1D), cilia elongation (Fig. 2D, 2E), and formation (Fig. 2D, 2F). We next applied this approach also to NIH3T3 cells, with very similar results - LGK974 inhibited WNT signaling (Fig 2G, 2H), but it failed to show any effect on cilia length, in contrast to CytoD treatment (Fig. 2I, 2J). We noted the CytoD treatment in NIH3T3 did not increase the cilia numbers (Fig 2K), possibly due to high basal ciliation rate of NIH3T3 compared to RPE-1. In sum, these data imply that signaling mediated by endogenous WNT ligands is not required for primary ciliogenesis.

**Figure 2:**
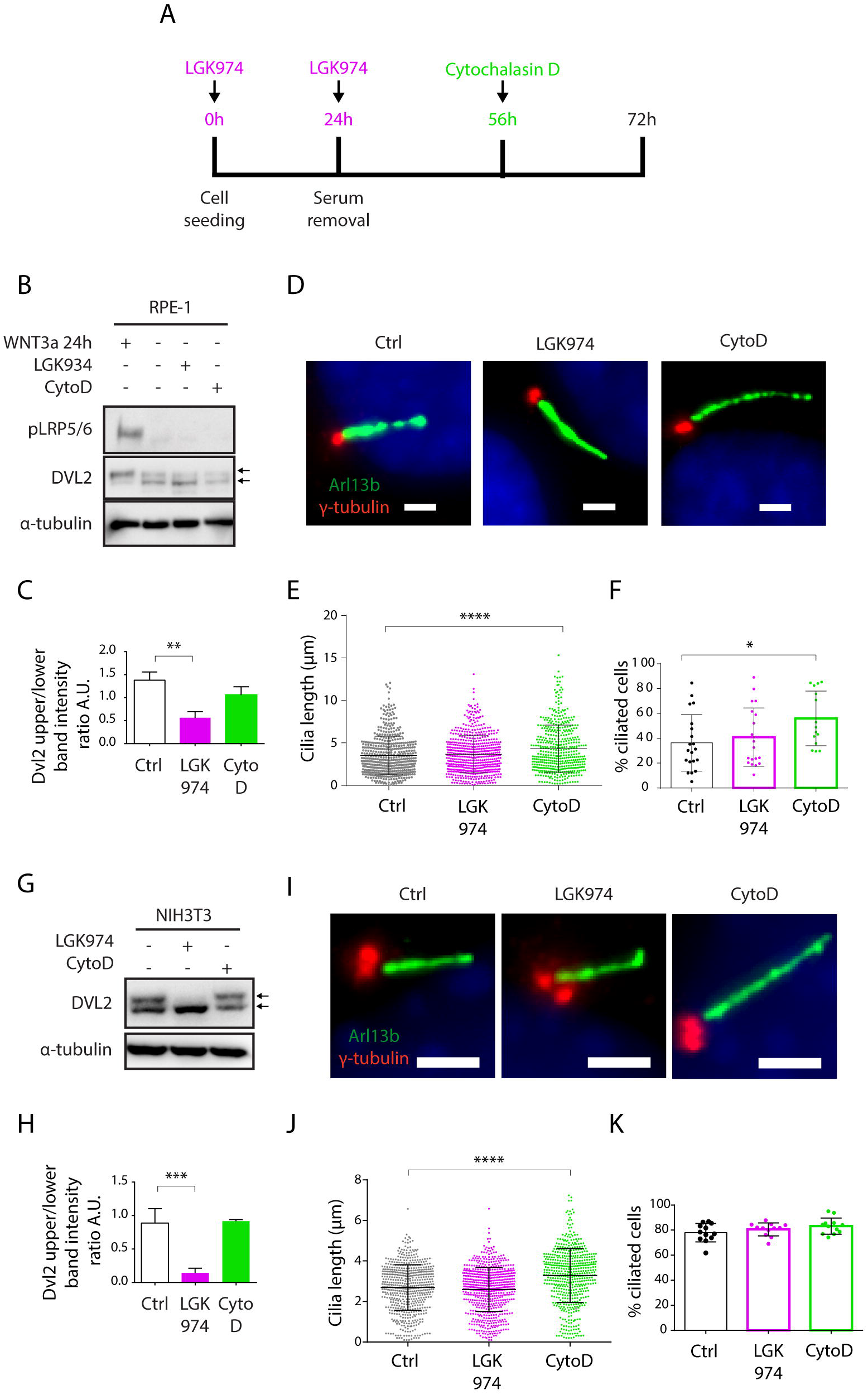
Inhibition of WNT secretion has no effect on ciliogenesis or cilia length. **(A)** Experimental scheme illustrating the time points of LGK974 (Purple) or CytoD (Green) treatments. **(B)** Western blot analysis of RPE-1 treated with LGK974 or CytoD. WNT3a was used as positive control to activate WNT/β-catenin pathway. Note that DVL2 shift (upper to lower band intensity ratio) following the LGK974 treatment was decreased over the control cells, as quantified in **(C)** n=3. **(D**) Representative images of RPE-1 cells following the indicated treatment, stained for Arl13b (green) and γ-tubulin (red). Scale bar 2μm. DAPI (blue) was used to counter stain nuclei. Quantification of the cilia length **(E)** and the percentage of cells with Arl13+ cilium **(F).** Each dot indicates either length of a single primary cilium (E) or percentage of ciliated cells in one image frame (F) n≥3. **(G)** Western blot analysis NIH3T3 treated with LGK974 or CytoD. **(H)** Quantification of DVL2 band intensities (upper to lower band intensity ratio) n=3. **(I)** Representative images of NIH3T3 cells following treatment with LGK974 or CytoD, stained for Arl13b (green) and γ-tubulin (red). Scale bar 2μm. DAPI (blue) was used to counter stain nuclei. Quantification of the cilia length **(J)** and the percentage of cells with Arl13+ cilium **(K).** Each dot indicates either length of a single primary cilium (J) or percentage of ciliated cells in one image frame (K) n=3.

### Genetic ablation of WNT/β-catenin pathway does not alter primary ciliogenesis

To corroborate our findings, we established a panel of HEK293 cells devoid of critical components of WNT signaling pathways. To specifically block the course of WNT/β-catenin pathway we used LRP5/6 double knock out HEK293 cells, to block the course of any WNT signaling pathway we used DVL1/2/3 triple knock out HEK293 cells (Paclíková et al. 2017) and to overactivate WNT/β-catenin pathway we used AXIN1/2 double knock out HEK293 cells.

First, we have verified successful disruption of LRP5 gene by sequencing (Suppl. Fig. 2A), and lack of LRP6 and pLRP5/6 signals in LRP5/6 null cells by western blot (Suppl. Fig. 2B, Fig. 3A) Furthermore, we confirmed these cells cannot activate WNT/β-catenin signaling (Suppl. Fig 2C). Similarly, we confirmed disruption of AXIN1 and AXIN2 genes in AXIN1/2 dKO by sequencing (Suppl. Fig. 2D), and lack of AXIN1 by western blot (Suppl. Fig. 2E). In addition, we observed that loss of AXIN1/2 function leads to excessive ABC accumulation (Fig. 3A) and in turn to overactivation of WNT/β-catenin signaling in AXIN1/2dKO cells (Suppl. Fig 2F), as expected.

**Figure 3:**
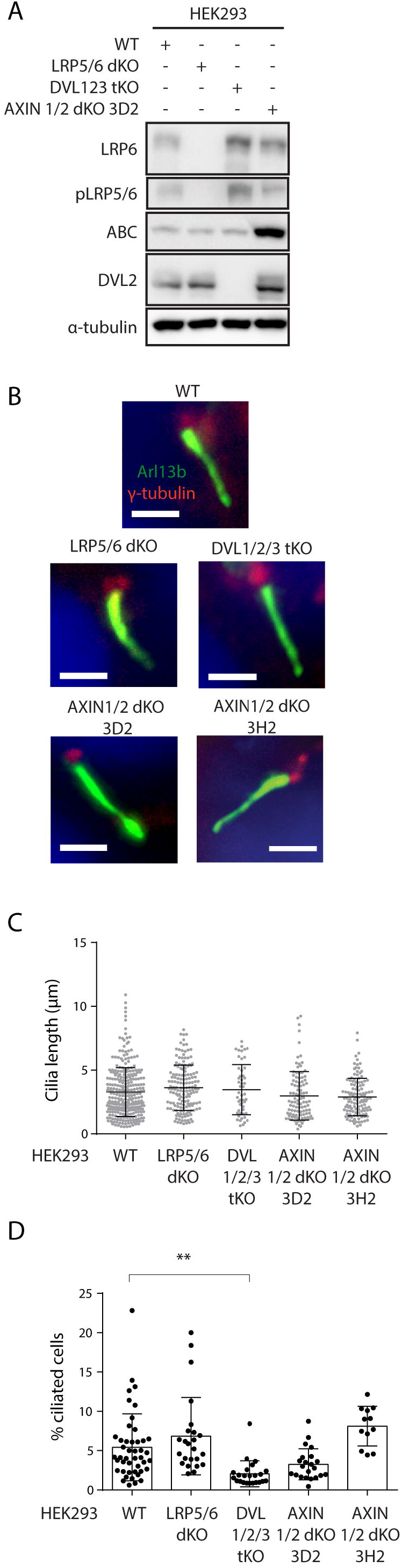
Ablation of WNT β-catenin pathway does not alter primary ciliogenesis. **(A)** Western blot analysis of individual HEK293 KO cell lines using the indicated antibodies. Note that LRP5/6 dKO cells lack LRP6 and phospho LRP5/6 (pSer1493/pSer1490), DVL1/2/3 tKO cell do not have detectable levels of DVL2. AXIN1/2 dKO cells shown elevated level of ABC. **(B-D)** HEK293 Cells were starved for 48h, stained for Arl13b (green), γ-tubulin (red), and DAPI (blue) and analyzed by IF microscopy. Representative images are shown in (B). Scale bar 2μm. Quantification of cilia length and percentage of ciliated cells is shown in (C) and (D), respectively Each dot indicates either length of a single primary cilium (C) or percentage of ciliated cells in one image (D). n=4.

Having characterized our model system, we examined cilia formation in those cells. Consistently with previous work, HEK293 cells form cilia less frequently than RPE-1 or NIH3T3 cells (Lancaster, Schroth, and Gleeson 2011; Bernatik et al. 2020). We were able to detect about 5% of cells with Arl13b+ primary cilium in WT HEK293. The percentage of ciliated cells, but not the cilia length, was reduced in DVL1/2/3 tKO cells (Fig. 3B, 3C, 3D). This observation is in agreement with the role of DVL and WNT/PCP pathway in the regulation of basal body positioning and ciliogenesis (Park et al. 2008; Shnitsar et al. 2015; Sampilo et al. 2018). Systemic activation of WNT/β-catenin pathway by AXIN1/2 removal produced a somewhat mixed result. Using AXIN1/2 dKO clone 3D2 we initially observed a non-significant negative trend on the cilia formation. However, this was not confirmed using an independent clone 3H2 (Fig. 3D). Importantly, the ablation of WNT/β-catenin pathway in LRP5/6 dKO cells had no effect on either the percentage of ciliated cells or cilia length (Fig. 3B, 3C, 3D), in agreement with our earlier observations based on pharmacological inhibition of endogenous WNT signaling in RPE-1 or NIH3T3. In sum, from these data we conclude that WNT/β-catenin signaling is not required for effective ciliogenesis.

## Discussion

Regulation of ciliogenesis is a complex process involving multiple factors directly or indirectly influencing cilia initiation and elongation. The regulators of cilium formation encompass a wide range of molecules such as components of centrioles, regulators of vesicular trafficking, intraflagellar transport proteins, membrane proteins and components of cytoskeleton (Wang and Dynlacht 2018; Conkar and Firat-Karalar 2020; Ishikawa and Marshall 2017; Seeley and Nachury 2010).

WNT3a is considered a prototypical “canonical” WNT ligand that activates WNT/β-catenin pathway (Willert et al. 2003). Moreover, WNT3a and hence the WNT/β-catenin pathway are well known for their mitogenic potential in many experimental systems (Niehrs and Acebron 2012). In addition, WNT/β-catenin pathway has been shown to act mainly during G2/M phase of the cell cycle (Davidson et al. 2009), while primary cilia form during G0/G1 and during the G2/M they disassemble (Ford et al. 2018; Rieder, Jensen, and Jensen 1979). Furthermore, mitogenic signals typically promote cilium disassembly (Rieder, Jensen, and Jensen 1979; Tucker, Pardee, and Fujiwara 1979; Pugacheva et al. 2007). From this perspective, the recently reported positive role of WNT3a and WNT/β-catenin signaling on primary cilia formation (Kyun et al. 2020) is counterintuitive and puzzling.

Principally, there are several important methodological differences between our work and the previous results (Kyun et al. 2020) which may account for the different outcomes. 1. In our experiments we activated the WNT/β-catenin pathway by recombinant WNT3a, in contrast to WNT3a conditioned medium often used in the previous study (Kyun et al. 2020). Thus, some of the reported effects of WNT3a conditioned medium may be a result of secondary effects. 2. We applied up to 24h stimulation by WNT3a to activate or 72h LGK974 to block the pathway, respectively. We cannot formally exclude that the longer WNT3a treatments used by Kyun et al., could account for the observed differences. However, we argue this seems unlikely, given that full activation of the WNT/β-catenin pathway or cilium formation typically happens within several hours following the proper stimuli (Wu, Chen, and Tang 2018; Pejskova et al. 2020; Lu et al. 2015; Bryja, Schulte, and Arenas 2007; Naik and Piwnica-Worms 2007; Pitaval et al. 2010). In fact, prolonged WNT/β-catenin pathway stimulation increases a chance for indirect secondary effects. Indeed, WNT signaling has been shown to regulate expression of a number ligands from FGF (Shimokawa et al. 2003; Hendrix et al. 2006; Chamorro et al. 2005; Kratochwil et al. 2002; Barrow et al. 2003) or BMP (J. S. Kim et al. 2002; Baker, Beddington, and Harland 1999; Shu et al. 2005) families that might in turn affect ciliogenesis (Cibois et al. 2015; Komatsu, Kaartinen, and Mishina 2011; Bosakova et al. 2018; Neugebauer et al. 2009). 3. Finally, we visualized cilia by staining for Arl13b, a small GTPase from Arf/Arl-family highly enriched in the ciliary membrane (Caspary, Larkins, and Anderson 2007; Hori et al. 2008; Cantagrel et al. 2008; Duldulao, Lee, and Sun 2009; Cevik et al. 2010; Li et al. 2010). In the report by Kyun et al., acetylated α-tubulin antibody staining was used to assess the cilia length, thickness, and numbers. From this perspective, it is plausible some of the reported changes in cilia length or thickness in fact reflect changes in the acetylation of ciliary tubulin rather than changes in cilium size. That being said, there is an evidence that individual cilia differ significantly in the levels of tubulin post-translation modifications and the levels of tubulin modifications may dramatically change in response to the appropriate stimuli (He et al. 2018; Piperno, LeDizet, and Chang 1987; Berbari et al. 2013).

Our data show that while WNT3a consistently activates the WNT/β-catenin pathway it has no or only minor negative effects on ciliogenesis. Elevated β-catenin levels following APC ablation have been related to reduced ciliogenesis and cell cycle defects in the developing cortex in mice (Nakagawa et al. 2017). We detected modest decrease in the percentage of ciliated NIH3T3 cells following WNT3a induced β-catenin accumulation. We speculate we did not observe comparable negative effect on cilia following the WNT/β-catenin pathway activation after AXIN1/2 loss due to abnormal cell cycle regulation in HEK293, which hampers detection of relativelly subtle deviations in their cell cycle progression (Stepanenko and Dmitrenko 2015; Löber, Lenz-Stöppler, and Dobbelstein 2002). These data are in contrast to Kyun et al., where accumulation of β-catenin by WNT3a conditioned medium treatment or by expression of S45A nondegradable oncogenic mutant variant of β-catenin (C. Liu et al. 2002) facilitates ciliogenesis.

In sum, we found no evidence that endogenous WNT/β-catenin signaling, while ablated either pharmacologically in RPE-1 or NIH3T3 by LGK974, or genetically by removal of LRP5/6 in HEK293, is required for primary cilia to form. Our findings presented in this article challenge some of the published evidence and argue against positive role of WNT3a or WNT/β-catenin pathway in ciliogenesis or cilia length regulation.

## Acknowledgement

We acknowledge the core facility CELLIM supported by the Czech-BioImaging large RI project (LM2018129 funded by MEYS CR) for their support with obtaining scientific data presented in this paper. hCas9(BB)-2A-GFP was a gift from George Church (Addgene plasmid #41815), pU6-(BbsI)_CBh-Cas9-T2A-mCherry was a gift from Ralf Kuehn (Addgene plasmid # 64324).

The work was supported by grants from the Czech Science Foundation (19-05244S) and the Swiss National Science Foundation (IZ11Z0_166533) to LC. OB was supported by funds from the Faculty of Medicine MU to junior researcher (Ondrej Bernatik, ROZV/28/LF/2020). VB was supported from European Structural and Investment Funds, Operational Programme Research, Development and Education – “Preclinical Progression of New Organic Compounds with Targeted Biological Activity” (Preclinprogress) - CZ.02.1.01/0.0/0.0/16_025/0007381.

## Notes

### Competing Interest Statement

The authors have declared no competing interest.

