## Supplementary material for "Primary cilia formation does not rely on WNT/β-catenin signaling": Suppl. Fig.1 and Fig.2

Supplementary Figure 1

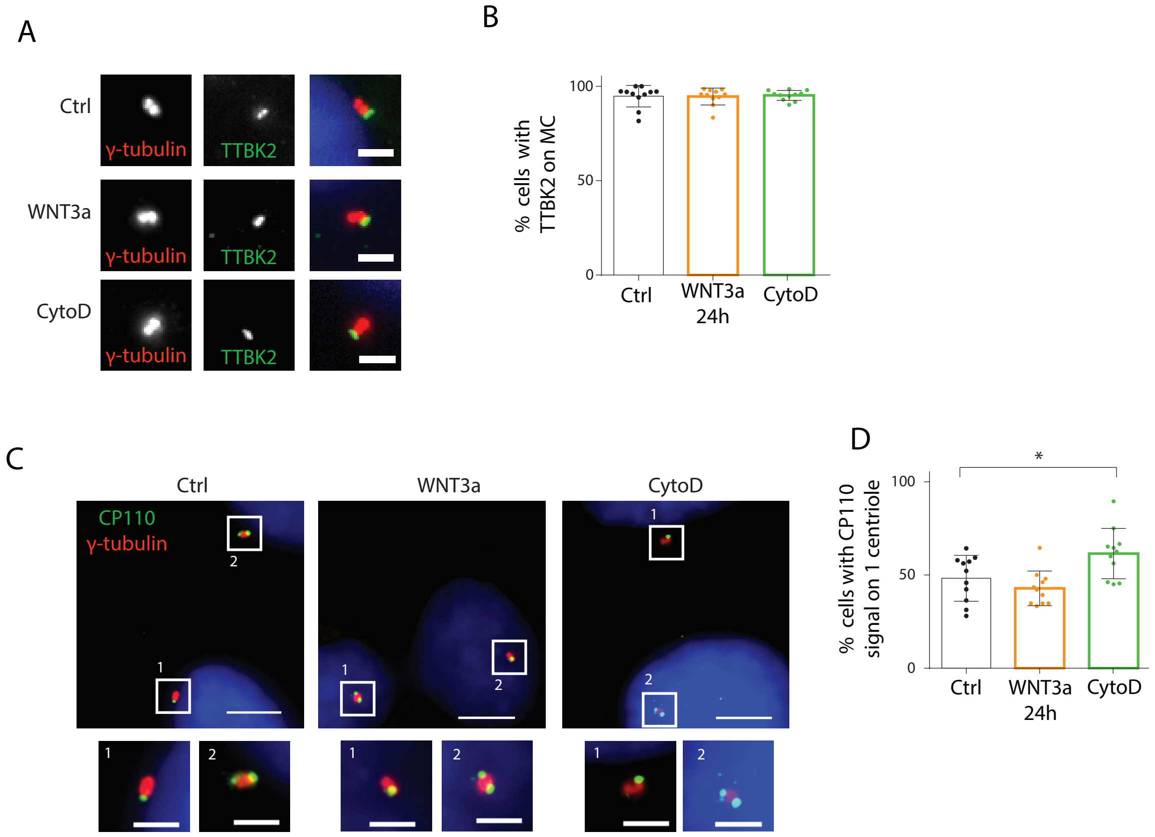


**Supplementary figure 1: WNT3a does not affect cilia initiation in RPE-1**
RPE-1 cells were treated by either WNT3a or CytoD (see Fig.1A for details) and stained by the indicated antibodies. **(A)** Representative images of IF staining for TTBK2 (green) and γ-tubulin (red), DAPI (Blue) was used to counter stain nuclei, Scale bar 2µm. The quantification of a percentage of cells with TTBK2 signal at MC (one dot represents single image) is shown in **(B)** n=3. **(C)** Representative images of staining for centriole distal end protein CP110 (green) and γ-tubulin (red). DAPI (Blue) was used to counter stain nuclei. White rectangles indicate centrosomes with CP110 signal on either one centriole (1) of both centrioles (2), the insets are enlarged below. The effects of WNT3a or CytoD are quantified in **(D)** each dot represents a percentage of cells with CP110 signal present only on one of the two centrioles, n=3. Scale bar 5µm.

Supplementary Figure 2

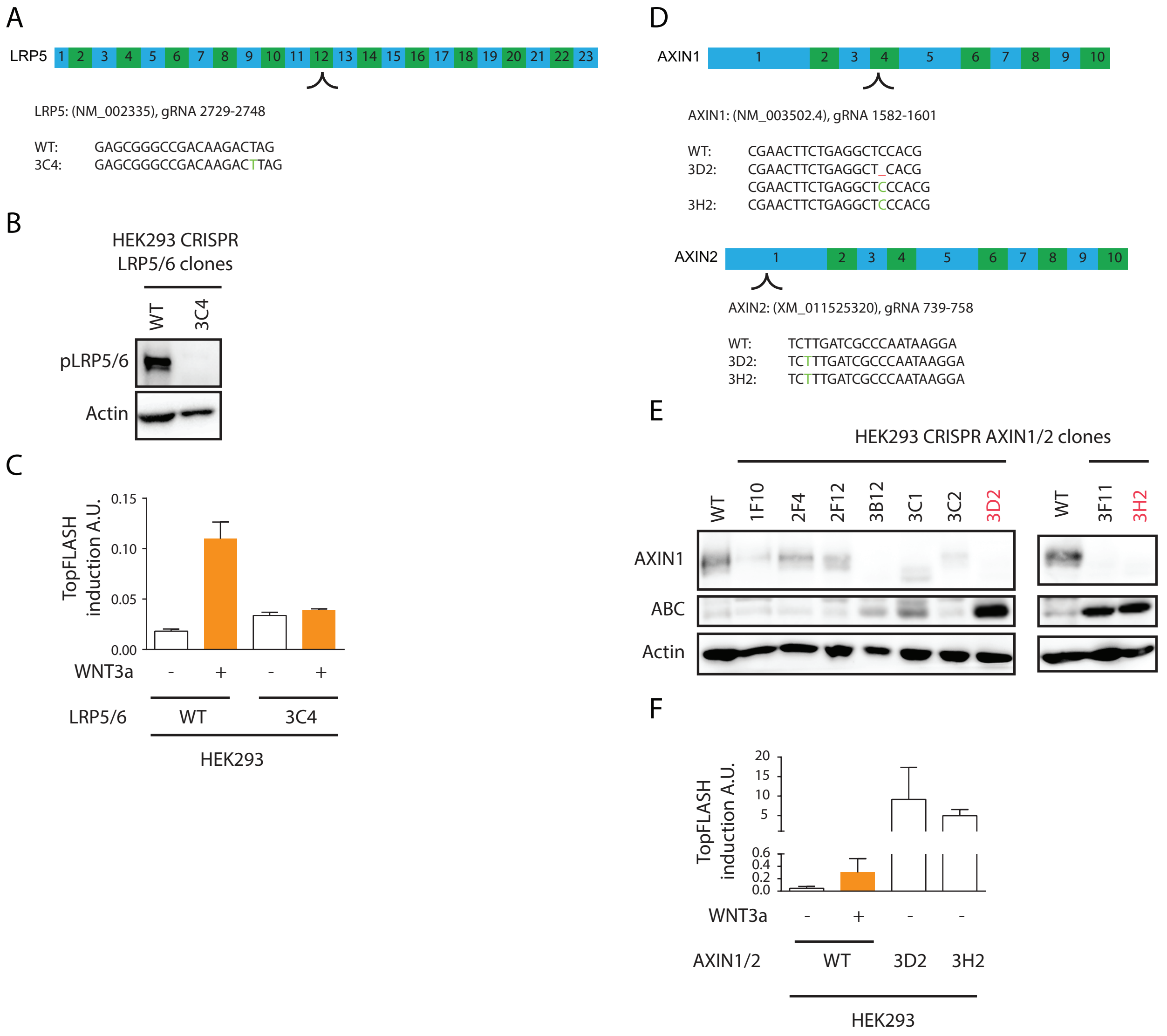


**Supplementary figure 2: Verification of LRP5/6 and AXIN1/2 knock-out cells**
**(A)** Scheme of LRP5 editing by CRISPR-Cas9. Individual exons are coded in blue and green. The used gRNA targets exon 12. Transcript variants unique identifier and exact position of gRNA are specified below. Cas9-edited sequence of LRP5 is indicated, the inserted “T” is highlighted in green. **(B)** Western blot analysis demonstrating the lack of phospho LRP5/6 signal in HEK293 clone 3C4 (LRP5/6 DKO). **(C)** Luciferase reporter assay showing that HEK293 clone 3C4 cannot activate WNT/β-catenin pathway. HEK293 WT and LRP5/6 dKO clone 3C4 were treated by WNT3a for 24h and activity of WNT/β-catenin pathway was measured by TopFLASH dual luciferase assay, n=3. **(D)** Schematic view of AXIN1 and AXIN2 editing by CRISPR-Cas9. Individual exons are coded in blue and green. Arrow indicates targeted exons for the used gRNAs, transcript variants unique identifiers and exact position of gRNAs are indicated below. Cas9-edited sequence of AXIN1 and AXIN2, detected indels are visualized in green and red, respectively. **(E)** WB detection of AXIN1 expression in the analyzed clones. Clones 3D2 and 3H2 which lacked AXIN1 expression and showed the highest levels of active β-catenin (ABC) were selected for further use. **(F)** Luciferase reporter assay showing that HEK293 clones 3D2 and 3H2 have overactive WNT/β-catenin pathway. Activity of the WNT/β-catenin pathway was measured by TopFLASH dual luciferase assay. 24h WNT3a treatment of HEK293 WT was used as positive control. Note that the reporter activity in untreated AXIN1/2 dKO clones 3D2 and 3H2 is notably elevated even if compared to WNT3a-treated WT cells.
